# Targeting Viperin to the Mitochondrion Inhibits the Thiolase Activity of the Trifunctional Enzyme Complex

**DOI:** 10.1101/824425

**Authors:** Arti B. Dumbrepatil, Kelcie A. Zegalia, Keerthi Sajja, Robert T. Kennedy, E. Neil. G. Marsh

## Abstract

Understanding the mechanisms by which viruses evade host cell immune defenses is important for developing improved antiviral therapies. In an unusual twist, human cytomegalovirus (HCMV) co-opts the antiviral radical SAM enzyme, viperin (**V**irus **i**nhibitory **p**rotein, **e**ndoplasmic **r**eticulum-associated, **in**terferon-inducible), to enhance viral infectivity. This process involves translocation of viperin to the mitochondrion where it binds the β-subunit (HADHB) of the mitochondrial trifunctional enzyme complex that catalyzes the thiolysis of β-ketoacyl-CoA esters as part of fatty acid β-oxidation. We have investigated how the interaction between these two enzymes alters their activities and their effect on cellular ATP levels. Studies with purified enzymes demonstrated that viperin inhibits the thiolase activity of HADHB, but, unexpectedly, HADHB activates viperin to synthesize the antiviral nucleotide 3’-deoxy-3’,4’-didehydro-CTP. Enzyme activities were also measured in lysates prepared from transfected HEK 293T cells transiently expressing these enzymes. Mirroring the studies on purified enzymes, localizing viperin to the mitochondria decreased thiolase activity whereas co-expression of HADHB significantly increased viperin activity. Furthermore, targeting viperin to mitochondria also increased the rate at which HADHB was retro-translocated out of mitochondria and degraded, providing an additional mechanism for reducing HADHB activity. Targeting viperin to the mitochondria decreased cellular ATP levels by over 50 %, consistent with the enzyme disrupting fatty acid catabolism. These results provide biochemical insight into the mechanism by which HCMV subjugates viperin; they also provide a biochemical rational for viperin’s recently discovered role in regulating thermogenesis in adipose tissues.

## Introduction

Viruses and cells have co-evolved such that the increasingly sophisticated antiviral defenses of host cells have been met with equally ingenious mechanisms by which viruses evade or neutralize them (1–5). The innate immune system provides the primary line of defense against viruses and other pathogens, with type 1 interferon (IFN) mediating the induction of over 300 IFN-stimulated genes that collectively act to disrupt viral replication and transmission (6–9). Viruses have evolved multiple strategies to evade various components of the innate immune system, which often appear to be virus-specific: these include binding cellular signaling proteins to prevent the expression of interferon stimulating genes (ISGs); proteolytic degradation or deubiquitination of cellular proteins by virally encoded proteases and deubiquitinases; and sequestering viral nucleic acids within cellular membranes (1,3,5,6,10,11). A better understanding of the diverse mechanisms by which viruses evade the immune system promises to lead to more effective antiviral drugs.

Here we have investigated one of the more unusual examples of a virus co-opting the cellular antiviral machinery to enhance infectivity. This involves human cytomegalovirus (HCMV), which co-opts the antiviral protein, viperin (**V**irus **i**nhibitory **p**rotein, **e**ndoplasmic **r**eticulum-associated, **in**terferon-inducible; also denoted as RSAD2 or Cig5 in humans), to facilitate the release of the virus from the cell (12,13).

Viperin appears to be conserved in all animals (12,14,15). Viperin-like enzymes have also been reported in fungi and archaea, indicating that it is an ancient component of the antiviral response (16,17). Viperin is a radical SAM enzyme, (18) one of only 8 annotated in the human genome (19). Radical SAM enzymes are predominately found in microbes where they catalyze a remarkably wide range of reactions that are initiated by the reductive cleavage of SAM to produce adenosyl radical (20–26). Viperin was recently shown to catalyze the formation of 3’-deoxy-3’,4’-didehydro-CTP (ddhCTP) through the dehydration of CTP by a radical mechanism (27). This novel nucleotide acts as a chain-terminating inhibitor of viral RNA-dependent RNA polymerases that are essential to the replication of flaviviruses (27).

Although the enzymatic activity of viperin explains its antiviral activity against dsRNA viruses such as flaviviruses, viperin exerts antiviral activity against a much wider range of both RNA and DNA viruses (14). It is also involved in regulating innate immune signaling (28) and lipid metabolism (29–31). Intriguingly, very recent studies on mice in which viperin was knocked out have implicated the enzyme in regulating thermogenesis in adipose tissues through regulation of fatty acid b-oxidation (32).

These varied aspects of viperin’s activity appear to result from interactions with a wide variety of other cellular and viral proteins. For example, we recently investigated viperin’s role in innate immune signaling through the Toll-like receptor-7 and 9 (TLR7/9) signaling pathways (33). We found that viperin mediated the formation of a complex between interleukin-1 receptor-associated kinase-1 (IRAK1) and the E3 ubiquitin ligase TNF receptor-associated factor 6 (TRAF6) and that this complex was necessary to observe Lys-63–linked polyubiquitination of IRAK1 by TRAF6, which is a key step in the signaling pathway. However, although IRAK1 ubiquitination appeared to depend on structural changes induced by SAM binding to viperin, it did not require catalytically active viperin (33).

Previous studies have shown that, perhaps counterintuitively, HCMV directly induces expression of viperin upon infection (12,13). Viperin is normally associated with the cytosolic face of the endoplasmic reticulum (ER), but the virally-encoded protein mitochondrial inhibitor of apoptosis (vMIA) binds to viperin and causes it to be targeted to the mitochondrion (Fig. 1) (13,34). In the mitochondrion, viperin binds to HADHB, the b-subunit of the mitochondrial trifunctional protein, (35) a multienzyme complex that catalyzes the last 3 steps in the β-oxidation pathway of fatty acids. Targeting viperin to the mitochondrion disrupts the cellular pools of ATP and NADH that, in turn, appears to disrupt the actin cytoskeleton (12,13). This disruption is hypothesized to aid newly synthesized viruses in escaping from the cell.

**Figure 1.**
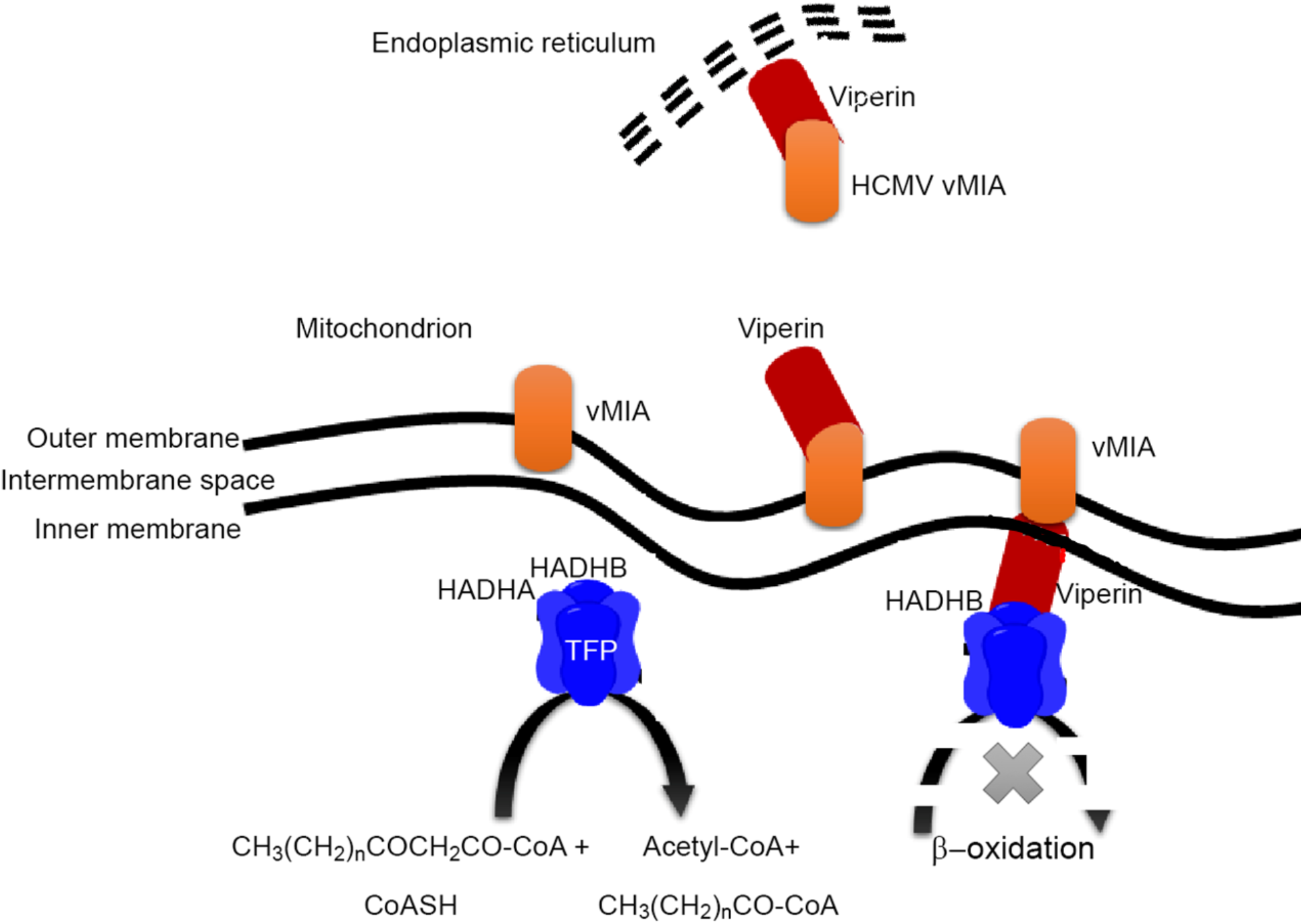
Human cytomegalovirus recruits viperin to the mitochondrion. The viral protein vMIA binds to viperin at the ER membrane and causes it to be translocated into the matrix of the mitochondrion. In the mitochondrion viperin is hypothesized to inhibits the thiolase activity of b-subunit of the mitochondrial trifunctional protein, preventing fatty acid oxidation from occurring.

However, the functional significance of viperin binding to HADHB remains unclear. Therefore, we have investigated the interaction between HADHB and viperin using both purified proteins *in vitro* and in transiently transfected HEK 293T cell lines that express viperin tagged with a mitochondrial localization signal (MLS). Our results provide support for the hypothesis that HCMV co-opts viperin to inhibit HADHB and thereby facilitate the escape of viral particles from the cell. They also throw light on the recent and intriguing observation that viperin is involved in regulating thermogenesis in murine adipose tissue (32).

## Results

### Viperin inhibits the thiolase activity of HADHB

The downstream effects on cellular physiology that result from vMIA translocating viperin to the mitochondrion have been attributed to inhibition of the fatty acid b-oxidation pathway (13). To test this hypothesis, we investigated what effect viperin had on the activity of HADHB *in vitro*. HADHB catalyzes the last step in b-oxidation, which is the thiolytic cleavage of the b-ketoacyl-CoA ester by CoASH to produce acetyl-CoA and the corresponding N-2-acyl-CoA ester (35). This reaction may be conveniently followed by measuring the decrease in absorbance at 303 nm due to the coordination of Mg^2+^ to the b-ketoacyl-CoA (36).

These experiments used a viperin construct lacking the first 50 residues corresponding to the N-terminal amphipathic helix, termed ΔN-viperin (29), as it has not proved possible to recombinantly express the full-length enzyme in active form. We also constructed an inactive mutant, ΔN-viperinC83A, in which one of the cysteine residues ligating the iron-sulfur cluster was mutated to alanine. Both ΔN-viperin and ΔN-viperinC83A proteins were expressed and purified from *E. coli* as described in the experimental section. Recombinantly expressed and purified HADHB was purchased from a commercial supplier (AdooQ^®^ Bioscience, Fig. S1c). Prior to experiments, the iron-sulfur cluster of viperin was reconstituted under anaerobic conditions to provide active holo-enzyme, as described previously (Fig. S2a) (29).

To investigate viperin’s effect on HADHB thiolase activity, assays were conducted under anaerobic conditions due to the air sensitivity of viperin. Reactions were performed at 22 °C and contained 100 μM acetoacetyl-CoA, 100 μM CoASH and 10 nM HADHB. Prior to assay HADHB was incubated with various concentrations of ΔN-viperin for 50 min. In the absence of viperin the apparent k_cat_ of HADHB was ~ 2 s^-1^ under the conditions of the assay. As the concentration of viperin was increased in the assay, the thiolase activity of HADHB was progressively inhibited, with the activity of HADHB declining to a minimum of 15 – 20 % of the initial activity at a 1:1 ratio of HADHB to viperin (Fig. 2a, Fig. S3). Interestingly, increasing the concentration of viperin did not result in any further decrease in HADHB activity, suggesting viperin does not act as a simple competitive inhibitor. In contrast, pre-incubation with ΔN-viperinC83A had no significant effect on HADHB activity (Fig. S4), although as discussed below the mutant still bound HADHB.

**Figure 2.**
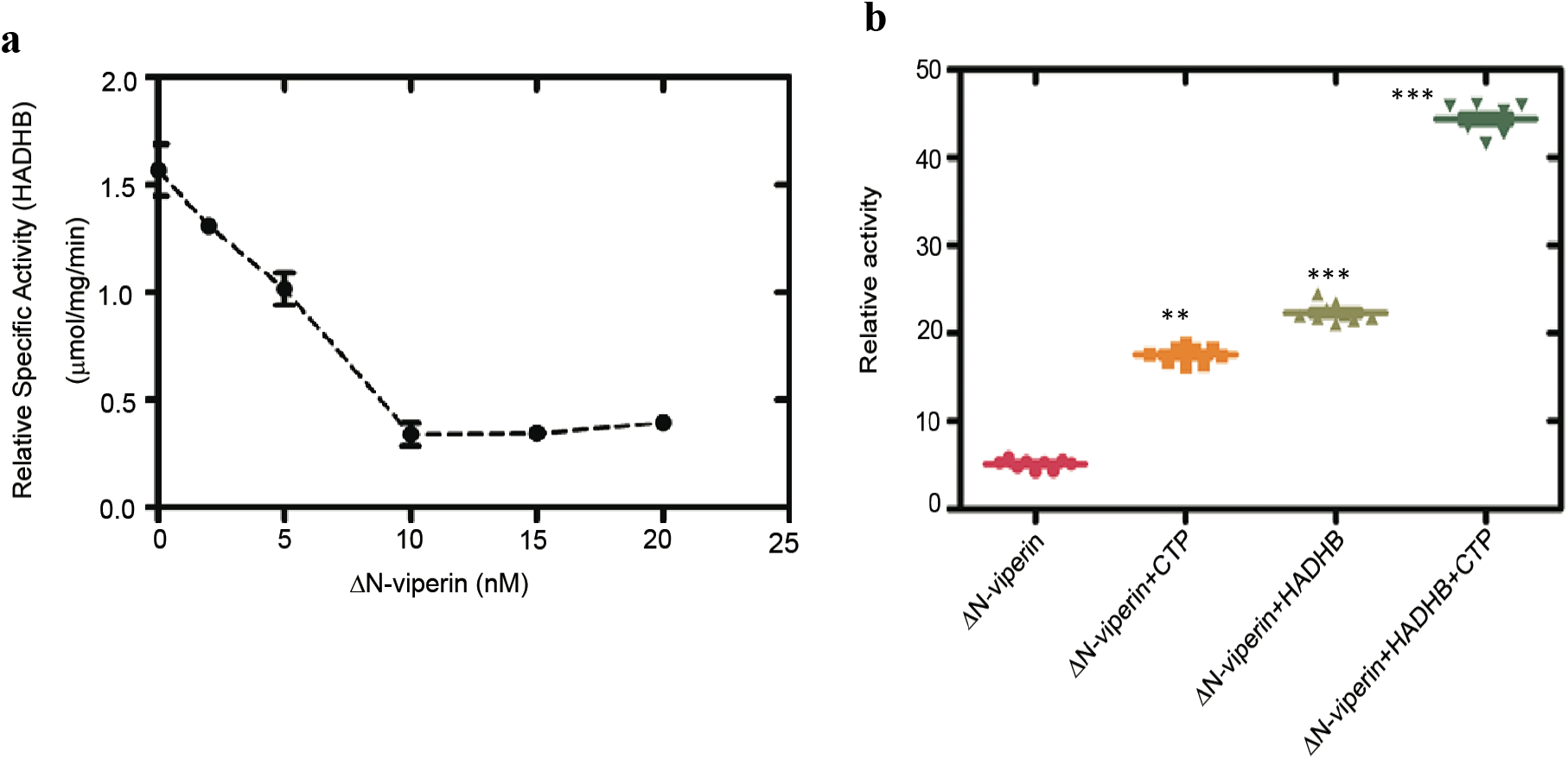
*In vitro* activity of HADHB and viperin. **a** The thiolase activity of purified, recombinant HADHB was assayed in the presence of increasing concentrations of ΔN-viperin. Maximal inhibition was observed at equimolar concentrations of the two enzymes **b** SAM cleavage activity of purified, recombinant ΔN-viperin assayed in the absence or presence of HADHB and/or the co-substrate 100 μM CTP. Both the uncoupled rate of SAM cleavage and the rate of SAM cleavage in the presence of CTP are significantly accelerated by HADHB. Values are presented as mean +/- S.E.M (n=6) with ** indicating p < 0.005 and *** indicating p < 0.001(Student’s t-test for independent samples).

### HADHB increases the activity of viperin

Previously we have shown that the enzymatic activity of viperin increases significantly when it forms a complex with IRAK1and TRAF6, as part of the TLR7/9 innate immune signaling pathway (33). This observation prompted us to investigate whether HADHB might similarly activate viperin (Fig 2b). Assays were conducted at 22 °C under anaerobic conditions as described in the experimental section. After 1 h, the reactions were quenched and the amount of 5’-dA formed was measured by liquid chromatography-mass spectrometry (LC-MS). In the absence of HADHB, the apparent *k*_cat_ of viperin was 2.2 ± 0.1 h^-1^, which is similar to values measured previously (27). However, in the presence of HADHB, the apparent *k*_cat_ of viperin increased by about 2.5-fold to 4.4 ± 0.1 h^-1^ (Table S1). Activation was specific to HADHB as no increase in activity was observed for viperin in the presence of proteins such as bovine serum albumin (Fig. S2b), which are often used to stabilize enzymes. It is interesting that although IRAK1/TRAF6 and HADHB perform very different functions, both activate viperin to catalyze the formation of ddhCTP, providing further support for the idea that protein-protein interactions play an important role in activating viperin.

### Mitochondrially targeting viperin results in HADHB degradation by the proteasomal pathway

Having established that ΔN-viperin inhibits HADHB *in vitro*, we extended our studies to examine how re-localization of viperin to the mitochondrion, as occurs in HCMV infection, may affect HADHB activity. Viperin is normally targeted to the membrane of the endoplasmic reticulum (ER) through its N-terminal amphipathic domain. Therefore, to redirect viperin to the mitochondrion this domain was replaced by a mitochondrial localization sequence (MLS). Residues 1–42, comprising the ER membrane-localizing domain of WT viperin were replaced by the MLS of the human mitochondrial protein Tom70 (residues 35–68), with this construct being designated MLS-viperin, as described previously (13).

We first established that HADHB and viperin interact within the mitochondrion by conducting pull-down experiments using transiently expressed proteins co-transfected in HEK 293T cells (Fig. 3a, Fig. S1a-d). Myc-tagged HADHB, which is naturally translocated into the mitochondrion, was co-transfected with either WT-viperin (which localizes to the ER membrane) or MLS-viperin. These experiments found that only MLS-viperin was co-precipitated with HADHB, consistent with viperin and HADHB forming a complex in the mitochondrial matrix. Fluorescence microscopy of fixed cells that had been transfected with MLS-viperin and stained with both MitoTracker-640, a mitochondrion-staining dye, and immuno-stained for viperin further confirmed that MLS-viperin co-localized with mitochondria (Fig. S1e). To further establish the interaction, we performed immunoprecipitation experiments with purified proteins. Both ΔN-viperin and ΔN-viperinC83A were efficiently pulled down by HADHB (Fig. 3b), confirming the interaction does not depend on the Fe-S cluster of viperin.

**Figure 3.**
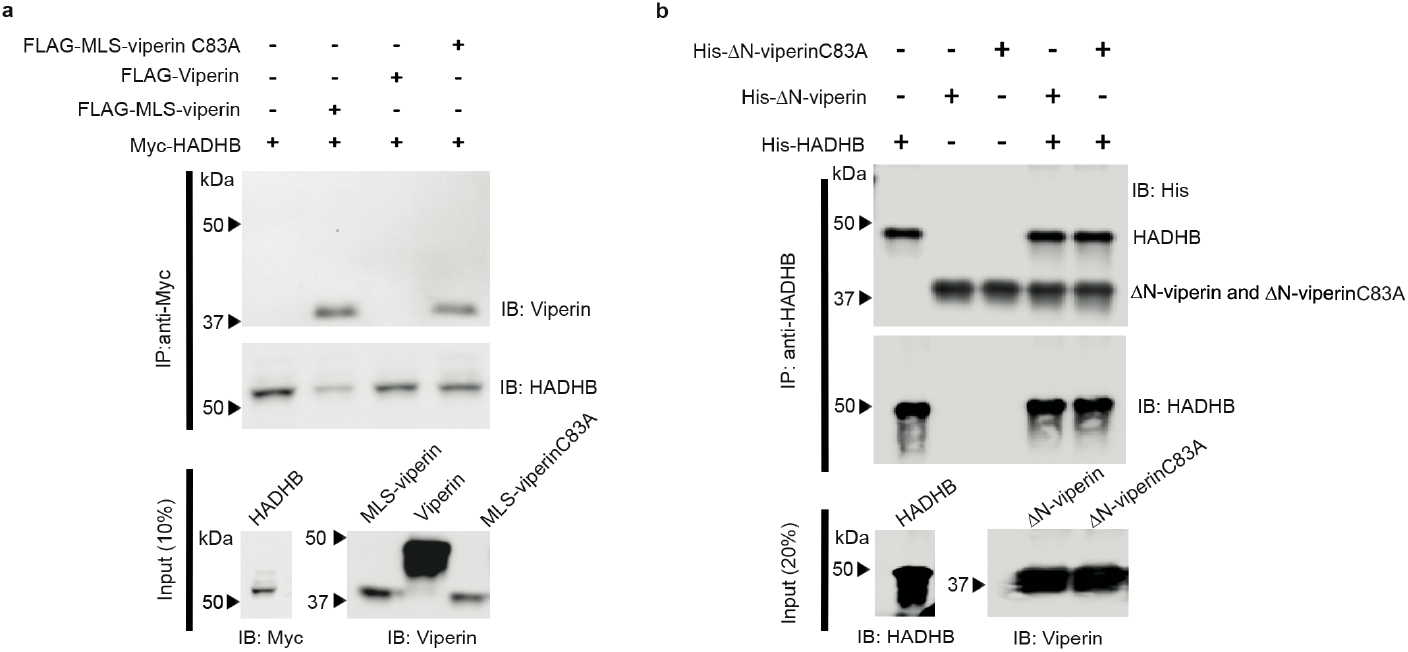
Viperin binds HADHB when targeted to the mitochondrion. **a** Immuno-tagged genes were transfected into HEK 293T cells and cell extracts were prepared 20 h post transfection. FLAG-MLS-viperin or FLAG-MLS-viperinC83A cell extracts were mixed with Myc-HADHB cell extracts in a ratio of 1:1. Proteins were immunoprecipitated with anti-Myc (HADHB) antibodies and analyzed by immunoblotting with the indicated antibodies. Control experiments confirmed the specificity of the antibodies used for immunoprecipitation, see Fig. S1 d. Immunoprecipitation of MLS-viperin and MLS-viperinC83A indicate that they bind to HADHB whereas cytosolically expressed viperin does not bind to HADHB. Mutations in the radical SAM domain do not affect ability of viperin to interact with HADHB. Representative blots are shown from two independent experiments. Cytosolic extracts were also immunoblotted to confirm expression levels of each individual protein of interest (10% input). **b** *In vitro* pull-down experiments were performed with recombinant proteins to examine the binding of HADHB to ΔN- viperin and ΔN-viperinC83A. Recombinant proteins purified from *E. coli* were immunoprecipitated with anti-HADHB antibody and the blots probed with anti-His antibody. As shown, ΔN-viperin and ΔN-viperinC83A were immunoprecipitated by anti-HADHB antibody specific to HADHB.

Viperin expression has been shown to be associated with the degradation of several cellular and viral proteins (14,29,37). Therefore, we next examined the effect of targeting viperin to the mitochondrion on the cellular levels of HADHB. The expression levels of HADHB were compared at 48 h following transfection of HEK 293T cells with Myc-tagged HADHB and either WT-viperin, MLS-viperin or MLS-viperin-C83A. In the latter viperin construct one of the three cysteines that coordinate the Fe4S4 cluster is mutated, thereby rendering the enzyme catalytically inactive.

In initial experiments, we found that co-expression of WT-viperin had no significant effect on the cellular concentration of HADHB; however, targeting viperin to the mitochondrion resulted in a significant decrease in HADHB levels (Fig. 4a). In contrast, co-expression of the inactive MLS-viperin-C83A mutant caused a much smaller decrease in HADHB levels. To investigate this observation in more detail, we studied HADHB levels as a function of time after co-transfection with MLS-viperin. The results show that MLS-viperin causes a time-dependent decrease in HADHB levels. However, in this experiment co-expression of MLS-viperin-C83A has no significant effect on HADHB levels (Fig. 4b). These results were confirmed when the experiment was repeated in the presence of cycloheximide (20 μM, CHX), to block *de-novo* protein synthesis (Fig. 4c).

**Figure 4.**
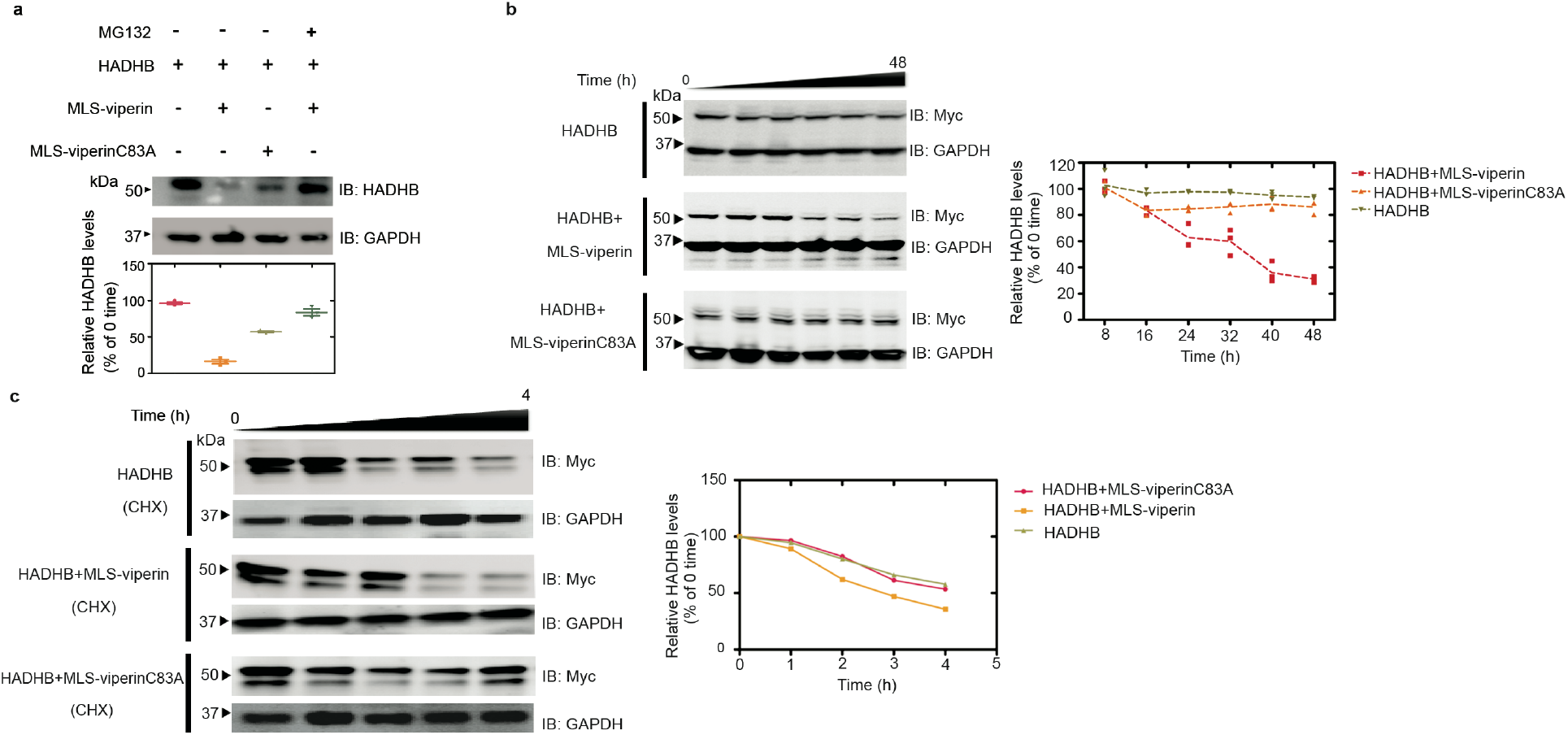
Targeting viperin to mitochondria results in degradation of HADHB. **a** Representative immunoblot of HEK293T cells transfected with HADHB, and either MLS-viperin or the inactive mutant MLS-viperinC83A, both in the presence and absence of the proteasomal inhibitor MG132. *Lower panel:* Quantification of HADHB levels determined by immunoblotting. **b** 48 h time course showing HADHB levels in cells transfected with HADHB, HADHB and MLS-viperin and HADHB and MLS-viperinC83A. **c** Cycloheximide chase followed by immunoblotting indicates that MLS-viperin accelerates HADHB degradation. HEK293T cells were transfected with HADHB, HADHB and MLS-viperin, or HADHB and MLS-viperinC83A and the amount of HADHB followed for 4 h after cycloheximide addition. HADHB levels are plotted as a function of time for each condition (t_o_ = 100%). The data are averages of 3 experiments with representative blots shown.

We also examined the effect of inhibiting the proteasomal degradation pathway on HADHB levels. Preliminary investigations established that HADHB levels are sensitive to the proteasomal inhibitor MG132, implying it is degraded by retro-translocation to the mitochondrial outer membrane followed proteolysis by the proteasomal pathway (Fig. 4a, Fig. S5). HEK 293T cells were co-transfected with HADHB and MLS-viperin and treated with MG132 (1 μM final concentration). The levels of HADHB in the cells were analyzed by immunoblotting. We observed that MG132 effectively counteracted the degradative effect of viperin on HADHB (Fig. 4a). This suggests that viperin reduces HADHB levels by increasing the rate of HADHB retro-translocation from the mitochondrion.

### Targeting viperin to the mitochondrion inhibits HADHB thiolase activity

To examine the effect of viperin on the activity of HADHB, cell lysates were prepared from HEK 293T cells that had been transfected with HADHB and co-transfected with WT-viperin, MLS-viperin or the MLS-viperin-C83A mutant. Cells were harvested 12 h after transfection, at which point HADHB levels were not yet significantly depleted by the action of viperin; cells transfected with empty vector served as a control. The thiolase activity of the lysates was measured using acetoacetyl-CoA and CoASH as substrates (36) as described in the experimental section.

The thiolase activity of cells transfected with HADHB alone was significantly higher compared to the background activity observed in cells transfected with an empty vector (Fig. 5a). After subtracting the background, the additional thiolase activity due to transfected HADHB was ~ 3.1 μM.min^-1^mg^-1^ of total protein. Lysates from cells co-transfected with HADHB and WT-viperin (which does not enter the mitochondria) exhibited similar levels of thiolase activity, 3.8 μM.min^-1^mg^-1^. However, in lysates prepared from cells co-transfected with HADHB and MLS-viperin the thiolase activity was significantly reduced to levels below that of the empty vector control. In contrast, lysates made from cells co-transfected with HADHB and MLS-viperinC83A exhibited very similar levels of thiolase activity to the HADHB-only lysates (Fig. 5a). These assays were carried out 12 h post-transfection because at this early time the HADHB levels did not vary greatly between the different conditions (Fig. S6). Therefore, differences in HADHB expression levels did not contribute significantly to the differences in thiolase activity.

**Figure 5.**
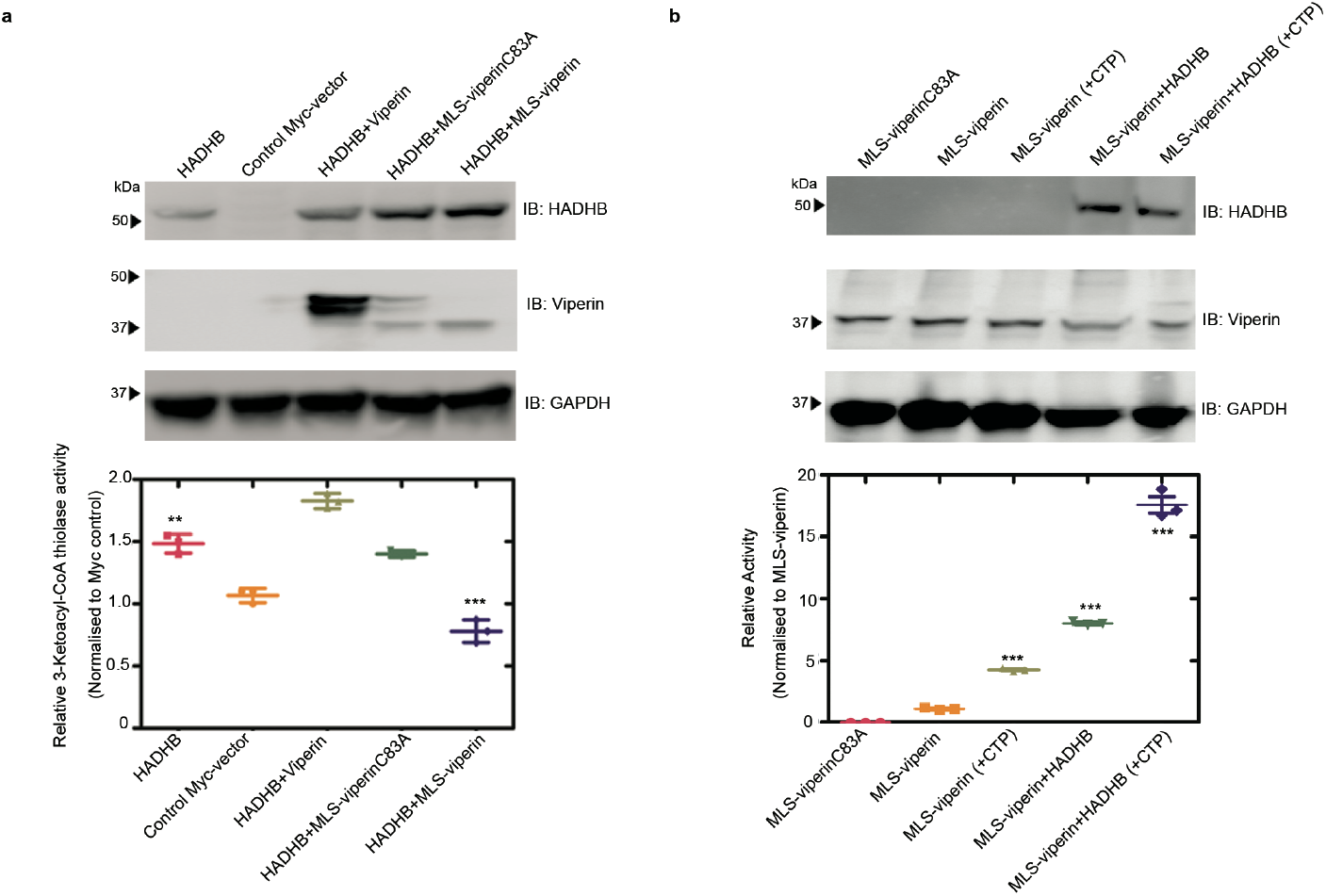
Effect of targeting viperin to mitochondria on the enzymatic activity of HADHB and viperin. **a** Cell lysates were prepared from HEK293T cells transfected with the indicated constructs and the thiolase activity of the lysate measured. *top panel*: expression of enzymes verified by immunoblotting; *bottom panel*: thiolase activity of the extracts relative to control. **b** Cell lysates were prepared from HEK293T cells transfected with the indicated constructs. Lysates were assayed anaerobically and contained 100 μM SAM and either no additional CTP or 100 μM CTP. *top panel*: expression of enzymes verified by immunoblotting; *bottom panel*: Activity is plotted relative to the MLS-viperin only condition (activity = 1) after normalizing for the viperin concentration. For details see the text. Values are presented as mean +/− S.E.M (n=6) with ** indicating p < 0.005 and *** indicating p < 0.001(Student’s t-test for independent samples).

### HADHB activates the radical SAM activity of viperin

The unexpected observation that, with purified enzymes *in vitro*, HADHB activates viperin prompted us to investigate whether HADHB exerted a similar effect on viperin in the context of the mitochondrial matrix (Fig 5b). HEK 293T cells were co-transfected with either MLS-viperin or MLS-viperinC83A and HADHB. 12 h post-transfection cell lysates were prepared under anaerobic conditions (due to the *in vitro* oxygen-sensitivity of viperin). The enzymatic activity of viperin was assayed under reducing conditions as described in the experimental section (33). The specific activity was quantified by measuring the amount of 5’-dA formed in 1 h, with the amount of viperin present in the cell extracts was quantified by immunoblotting, using methods described previously (33). Control experiments confirmed that the substitution of the N-terminal of viperin with the MLS sequence had no effect on viperin’s radical SAM activity (Fig. S4).

In cell lysates transfected with MLS-viperin alone, the specific activity of viperin, expressed as a turnover number, was 12.1 ± 0.5 h^-1^. Viperin activity increased significantly when HADHB was co-expressed in the cells by ~ 2-fold to give a turnover number of 27 ± 1 h^-1^. It appears, therefore, that HADHB activates viperin in the mitochondrion. We note that these turnover numbers are considerably higher than those measured with purified enzymes, but are similar to those we previously reported for viperin activation by TRAF6/IRAK1 (33). This discrepancy may, in part, reflect differences in the methods used to determine viperin concentration, but also suggests that when expressed in eukaryotic cells viperin is intrinsically more active. The availability of the endogenous iron-sulfur cluster-assembling machinery in eukaryotic cells may be one factor that contributes to the higher activity of viperin.

Interestingly, we also observed significant activation of viperin by HADHB in assays conducted in the absence of CTP. Under these conditions, lysates of cells transfected with only MLS-viperin exhibited a low level of activity, 1.6 ± 0.1 h^-1^, that may represent uncoupled reductive cleavage of SAM and/or reaction with endogenous CTP in the cell lysates (Fig. 5b). However, when co-expressed with HADHB, this “background” activity increased ~4-fold to 6.7 ± 0.3 h^-1^, an activity substantially higher than in lysates in which CTP was added but HADHB was absent (Table S2).

### Targeting viperin to the mitochondrion reduces cellular ATP levels

HADHB is a component of the multi-enzyme complex involved in the b-oxidation of fatty acids, which constitutes a major route for ATP production. Therefore, we examined the effect of targeting viperin to the mitochondrion on cellular ATP levels. We measured cellular ATP levels in transfected HEK 293T cells expressing HADHB and/or MLS-viperin, or MLS-viperinC83A, with an empty vector serving as a control. The results, summarized in Fig. 6, show that ATP levels were reduced significantly in cells expressing MLS-viperin, by over 50%. In contrast, ATP levels were only slightly reduced in cells expressing the inactive MLS-viperinC83A mutant. Interestingly, attempts to restore ATP levels by over-expressing HADHB with MLS-viperin did not result in an increase in ATP levels, suggesting that viperin is a potent inhibitor of HADHB. The reduction in cellular ATP levels is consistent with earlier studies (13) as well as our observation that the interaction of the mitochondrially-targeted viperin with HADHB reduces the β-thiolase activity and thus blocks b-oxidation.

**Figure 6.**
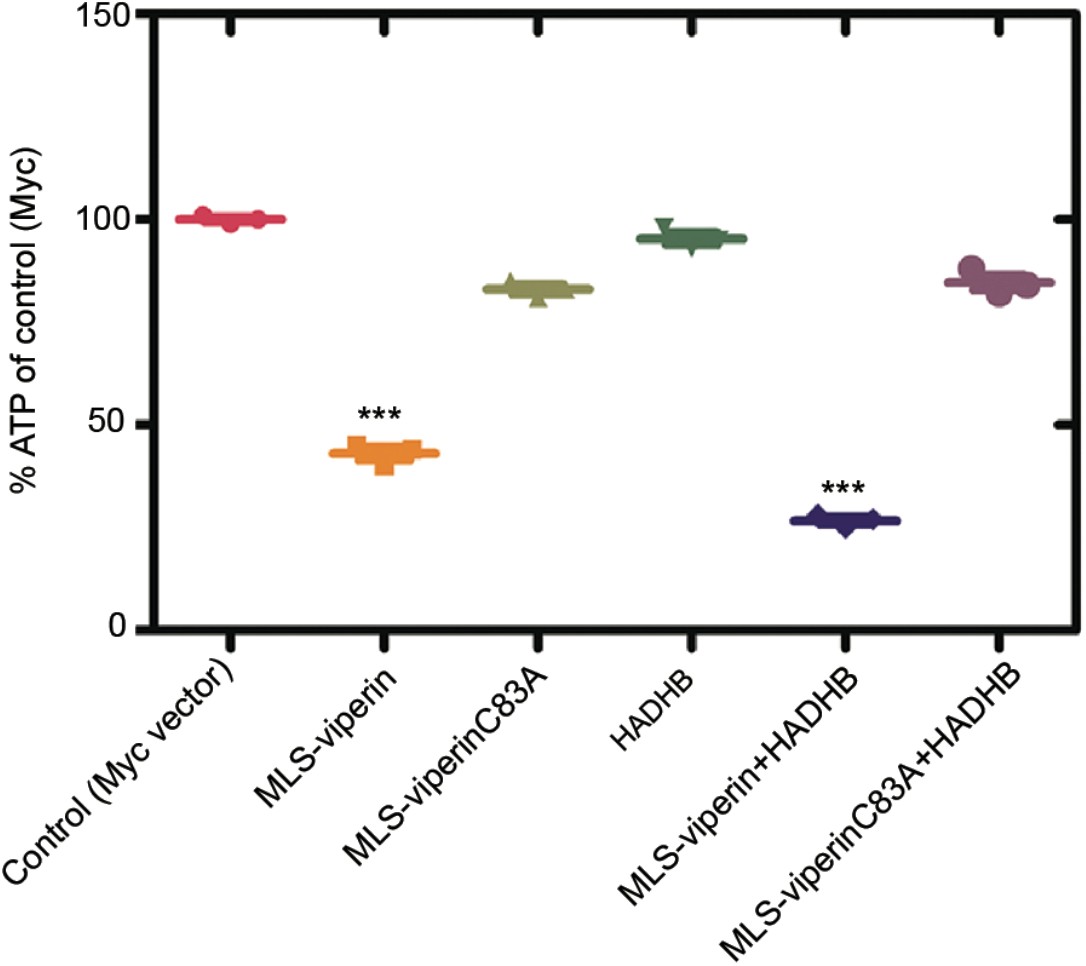
Targeting viperin to mitochondria results in a decrease in cellular ATP levels. HEK 293T cells were transfected with the indicated constructs and 24 h after transfection the cells harvested, and ATP concentrations determined. Results are plotted relative to the empty vector control = 100 %. Values are presented as mean +/− S.E.M (n=6) with *** indicating p < 0.001(Student’s t-test for independent samples).

## Discussion

There is considerable interest in the mechanisms by which viruses evade cellular defenses because a detailed understanding of such mechanisms is a prerequisite for developing new antiviral therapies. The co-opting of viperin by HCMV that leads to increased infectivity was first described by Cresswell and co-workers (13). They established that the viral protein vMIA is responsible for relocating viperin to the mitochondrion and that this results in a significant decrease in cellular ATP levels. Their investigations also identified HADHB as a target of viperin and on this basis they hypothesized that viperin inhibited fatty acid b-oxidation. They further hypothesized that the reduction in cellular ATP levels contributed to the observed disruption of the actin cytoskeleton, thereby facilitating the release of the virus from the cell. However, at the time of these studies, the biochemical function of viperin was unknown, which limited the conclusions that could be drawn regarding its effects on mitochondrial metabolism.

Through experiments with purified enzymes, we directly demonstrated that ΔN-viperin tightly binds to HADHB and inhibits its enzymatic activity, indicating that the two enzymes form a stoichiometric complex. This is consistent with the original observation that targeting viperin to the mitochondrion inhibits fatty acid oxidation and causes the down-stream effects on ATP levels and cytoskeleton integrity (13). Our results further suggest that vMIA is only responsible for transporting viperin into the mitochondrion but does not play a further role in mediating the inhibition of HADHB by viperin. Most interestingly, HADHB activity was not completely inhibited by ΔN-viperin, even when the enzyme was in 2-fold excess (Fig 2a). This suggests that viperin may regulate HADHB activity by an allosteric mechanism, rather than simply preventing the substrate from binding. It is also interesting that although the ΔN-viperinC83A mutant still bound to HADHB, it had no effect on activity. It therefore appears that the Fe-S cluster may play an important role in viperin’s regulation of HADHB.

Previous studies did not examine the effect of targeting viperin to the mitochondrion on the thiolase activity of HADHB. Here we show that co-expression of MLS-viperin with HADHB results in a significant decrease in thiolase activity in cell extracts that do not occur with WT-viperin. This decrease could be due to viperin reducing the cellular levels of HADHB, as appears to occur with other protein targets of the enzyme (29,30,37), or viperin could inhibit HADHB directly by binding to it. We found that co-expression of MLS-viperin with HADHB does result in a significant decrease in the cellular levels of HADHB, which could be reversed by blocking the proteasomal degradation pathway. This observation suggests that viperin causes HADHB to be retro-translocated to the outer mitochondrial membrane where it is subject to ubiquitination and subsequently degraded by the proteasome in a process known as outer mitochondrial membrane-associated degradation (38,39). However, details of the mechanism by which viperin binding to HADHB leads to its retro-translocation out of the mitochondrion remains to be elucidated. Overall, our data, summarized in Figs. 2, 4, and 5 indicate that viperin may regulate HADHB both by directly inhibiting it and through proteostasis by increasing its rate of degradation.

The original discovery that HCMV up-regulates viperin to enhance viral infectivity (13) was surprising because it appeared to rely on a seemingly adventitious association between viperin and HADHB. However, the recent finding that viperin is intrinsically expressed in mouse adipose tissue where it appears to regulate thermogenesis (32) suggests to us that viperin binding to HADHB likely serves a pre-existing regulatory function that the virus has evolved to exploit. In particular, the observation that mice lacking viperin exhibited increased heat production when fed a high-fat diet is consistent with viperin either inhibiting HADHB and/or reducing HADHB levels in the mitochondrion, as we have demonstrated, and thereby regulating b-oxidation. These observations suggest that the interaction between viperin and HADHB may provide a new target for anti-HCMV therapeutics and the regulation of thermogenesis in metabolic diseases.

The activating effect of HADHB on viperin is an interesting and unanticipated observation. We observed in both HEK 293T cell lysates and with purified proteins that HADHB activates the radical SAM activity of viperin by several-fold. Whether the production of ddhCTP in the mitochondrion also contributes to enhancing the infectivity of HCMV or plays a role in regulating thermogenesis is at this point unclear. ddhCTP has been shown to inhibit the replication of some RNA viruses by acting as a chain terminator during replication of the genome by the viral RNA-dependent RNA polymerases (27). Although ddhCTP does not appear to be incorporated by the nuclear RNA polymerases, the mitochondrial polymerase is structurally most closely related to bacteriophage T7 polymerase (40) and so might be susceptible to mis-incorporating ddhCTP. One intriguing possibility is that ddhCTP might act as a regulator of mitochondrial RNA transcription under non-pathological conditions by causing low levels of chain termination, if it were miss-incorporated. Under pathological conditions, such as HCMV infection, the production of ddhCTP might play a further role in enhancing virus infectivity by disrupting mitochondrial transcription more extensively.

## Experimental procedures

### Cell lines

The HEK293T cell line was obtained from ATCC.

### Antibodies

Rabbit polyclonal RSAD2/ Viperin antibody (11833-1-AP) was obtained from Protein Tech. Mouse monoclonal viperin antibody clone MaP (MABF106) was obtained from EMD Millipore Corporation. Rabbit polyclonal Myc-tag antibody (16286-1-AP) was obtained from Proteintech. (Mouse monoclonal HADHB(E-1) (sc-271495) was obtained from Santa Cruz. Goat anti-rabbit (170–6515) and anti-mouse (626520) Ig secondary Abs were purchased from BioRad and Life Technologies respectively. Rabbit polyclonal GAPDH (TAB1001) was purchased from Thermo Scientific and Mouse monoclonal GAPDH antibody (6C5) was obtained from EMD Millipore.

### Plasmids

HADHB (Myc-DDK-tagged)-Human hydroxyacyl-CoA dehydrogenase/3-ketoacyl-CoA thiolase/enoyl-CoA hydratase (trifunctional protein) plasmid (RC209853) was purchased from Origene (Rockville, MD, USA). MLS-viperin plasmid was a kind gift from Dr. Peter Cresswell, Yale School of Medicine, New Haven, CT 06520-8011. The MLS-viperinC83A mutant was constructed by site-directed mutations of the first cysteine residue to alanine (C83A) using the QuickChange Site-Directed Mutagenesis kit (Agilent). The following primer pair was used for introducing the change: 5’-CACTTCACTCGCCAGTGCAACTACAAATGCGGC-3’ and 5’-GCCGCATTTGTAGTTGCACTGGCGAGTGAAGTG-3’. Synthetic genes encoding human viperin (GenBank accession numbers AAL50053.1) were purchased from GenScript. pCMV-Myc plasmid (empty vector) (#631604) was purchased from Takara Biosciences.

### Recombinant human HADHB

Purified, recombinantly-expressed HADHB (AP4092) was purchased from AdooQ Bioscience LLC.

### Reagents

The sources of all other reagents have been described previously (29,33)

### Transfection

HEK293T cells were maintained in Dulbecco’s modified Eagle’s medium containing 10% fetal calf serum, 100 U/ml penicillin, 100 g/ml streptomycin, 2 mM L-glutamine. Transient transfections were carried out either using FuGENE^®^ HD (Promega) or PEI MAX - transfection grade linear polyethyleneimine hydrochloride (Polysciences Inc.) following the manufacturer’s instructions.

### Expression and Purification of recombinant truncated viperin (ΔN-Viperin /ΔN-ViperinC83A)

For *in vitro* assays of viperin, a truncated version of the enzyme (pET28b-ΔN-viperin)that lacked the N-terminal amphipathic helix was used. This enzyme, ΔN-viperin, was overexpressed and purified from *E. coli* as described previously (21, 29) The pET28b-ΔN-viperinC83A mutant was constructed by site-directed mutations of the first cysteine residue to alanine (C83A) using the QuickChange Site-Directed Mutagenesis kit (Agilent). The following primer pair was used for introducing the change: 5’-CACTTCACTCGCCAGTGCAACTACAAATGCGGC-3’ and 5’-GCCGCATTTGTAGTTGCACTGGCGAGTGAAGTG-3’. Expression, purification, and reconstitution conditions were similar to the wild type construct as described above.

### Assays of viperin activity using purified recombinant ΔN-viperin

Enzyme assay reactions were performed under anaerobic conditions in 10 mM Tris-Cl buffer pH 8.0, 300 mM NaCl, 10% glycerol and 5 mM sodium dithionite. Assays contained in a total volume of 100 μL: 10 nM ΔN-viperin, 100 μM SAM and 100 μM CTP. The assay was incubated for 60 min. at room temperature, after which time the reaction was stopped by heating at 95 °C for 10 min. The solution was chilled to 4 °C, and the precipitated proteins were removed by centrifugation at 14,000 rpm for 25 min. For assays with HADHB, HADHB was added at a concentration of 10 nM and other conditions were maintained the same. The 5’-dA generated during the reaction was then extracted with acetonitrile, as described previously (29,33). Samples were analyzed in triplicate by UPLC-tandem mass spectrometry as described previously (29,41).

### Assay of MLS-viperin activity in HEK 293T cell lysates

HEK 293T cells transfected with MLS-viperin, MLS-viperin1C and/or HADHB were harvested from one 10 cm diameter tissue culture plate each, resuspended in 400 μl of anoxic Tris-buffered saline (50 mM Tris-Cl, pH 7.6, 150 mM NaCl) containing 1% Triton X-100, sonicated within an anaerobic glove box (Coy Chamber), and centrifuged at 14,000 g for 20 min. Dithiothreitol (DTT; 5 mM) and dithionite (5 mM) were added to the cell lysate together with CTP (300 μM). The assay mixture was incubated at room temperature for 30 min prior to starting the reaction by the addition of SAM (200 μM). The assay was incubated for 50 min at room temperature, after which the reaction stopped by heating at 95 °C for 10 min. The solution was chilled to 4 °C, and the precipitated proteins were removed by centrifugation at 14,000 rpm for 25 min. The supernatant was then extracted with acetonitrile and the samples derivatized with benzoyl chloride as described previously (41) and analyzed in triplicate by UPLC-tandem mass spectrometry as detailed in the supporting information. For details of standard curve construction and calculations refer to (33). The viperin activity measurements reported here represent the average of at least three independent biological replicates.

### Immunoblotting and Immunoprecipitation Assays

Immunoprecipitation using the HEK 293T cell lysates was performed with anti-*Myc* tagged magnetic beads according to the manufacturers’ protocol, as detailed previously (32). Immunoprecipitation for *E.coli* purified proteins using Protein A beads was performed according to the manufacturers protocol. For immunoprecipitation, the ratio of suspension to packed gel volume was 2:1. Resin pre-equilibration was done as per the manufacturer’s protocol. The purified proteins, HADHB:ΔN-viperin / ΔN-viperinC83A were mixed in 1:1 ratio and incubated for 2 hours at 4°C with gentle rotation. Beads (attached to HADHB antibody) were pelleted by centrifugation at 5,000 × *g* for 30 seconds at 4°C and washed three times with washing buffer (50 mM Tris, pH 7.4,150 mM NaCl, 10% glycerol, 1 mM EDTA, 10 mM NaF, and 0.2 mM phenylmethylsulphonyl fluoride with protease inhibitor cocktail from Sigma). Immunocomplexes were eluted by boiling in SDS-PAGE sample buffer, separated by SDS-PAGE, and transferred to a PVDF membrane. Immunoblotting was performed using standard methods as described previously (33), with total protein concentrations determined by the BCA method and sample loading concentrations normalized accordingly. Blots were visualized, and band intensities quantified using a Bio-Rad ChemiDoc Touch imaging system, with GAPDH used as a control to compare relative expression levels. Quantitative measurements of protein expression levels reported here represent the average of at least three independent biological replicates. Quantitative measurements of protein expression levels reported here represent the average of at least three independent biological replicates.

### Assay of HADHB Thiolase activity using purified recombinant protein

Thiolase activity of HADHB was measured using acetoacetyl-CoA and CoASH as substrates and following the decrease in absorbance at 303 nm as described by Zhou et al (36). A typical assay contained 100 mM Tris-Cl pH 8.3, 25 mM MgCl2, 1 mM DTT, 100 μM CoASH, 100 μM acetoacetyl-CoA, and 10 nM HADHB complex in a 1 mL reaction volume. The reaction was started by adding HADHB and after 5 min incubation at room temperature the decrease in absorbance at 303 nm was monitored; under these conditions, the reaction rate remained linear for at least 30 min. The rate of reaction was calculated assuming an extinction coefficient of 16,900 cm^-1^ M^-1^ for the complex of acetoacetyl-CoA with Mg^2+^.

For thiolase activity measurements in the presence of viperin, the assay was performed with the following modifications. All the assay buffers were made anaerobic prior to the assay and the assay components assembled in anaerobic cuvettes in an anaerobic glove box (Coy Laboratory). Prior to addition to the assay mixture, HADHB (10 nM) and ΔN-viperin were mixed together at varying concentrations of ΔN-viperin (0, 2, 5, 10, 15, and 20 nM) under anaerobic conditions and incubated with gentle agitation at room temperature for 50 min. After adding the enzyme, the cuvettes were sealed to the atmosphere and removed from the anaerobic chamber and introduced into the spectrophotometer, with reaction rates being recorded 5 min after the addition of enzyme. For follow-up assays ΔN-viperin (10 nM) and HADHB (10 nM) were mixed together in equimolar concentrations and assayed as described above. For ΔN-viperinC83A assays similar conditions were followed. The thiolase activity reported here represents the average of at least three independent measurements.

### Assay of HADHB thiolase activity in HEK 293T cell lysates

Cell pellets were thawed on ice and were suspended in 300 μL lysis buffer (100 mM Tris-Cl pH 8.3, 2 mM b-mercaptoethanol, 200 mM NaCl, 0.5 mM EDTA, and 0.5 % Tween 20) containing Halt protease inhibitor cocktail. Cells were lysed by brief sonication. Cell debris were removed by centrifugation of the lysate at 14,000 rpm for 25 minutes at 4° C. The clarified lysates were used in thiolase assays, with the total protein concentration of the lysate determined using the BCA assay (Pierce™ BCA Protein Assay Kit, Thermo Scientific). A typical assay contained 10 μL cell lysate, 10 mM Tris-Cl, 25 mM MgCl2, 100 μM CoA, 1 mM DTT, and 100μM acetoacetyl-CoA in a 1 mL reaction volume. Thiolase activity was measured spectrophotometrically using the protocol described above. HADHB concentrations were determined by immunoblotting as described above. Thiolase activities were normalized to *Myc*-only control cell lysates. Measurements of thiolase activities reported here represent the average of at least three independent biological replicates.

### ATP assays

HEK 293T cells were transfected with indicated constructs and cellular ATP levels were measured with the ATP bioluminescence assay kit HS II (Roche) following the manufacturer’s instructions. To normalize the measured ATP levels, the total protein content of transfected cells was measured using the Bio-Rad protein assay kit. Serial dilutions of BSA were used as standards. Finally, ATP in cells transfected with indicated constructs was normalized to those in control cells transfected with vector alone.

### Statistical analyses

Results from all studies were compared with unpaired two-tailed Student’s *t*-test using GraphPad Prism 5 software. *P* values less than 0.05 were considered significant.

## Acknowledgements

We thank Prof. Peter Cresswell, Yale University for the kind gift of the MLS-viperin expression vector. This work was supported in part by NIH grants GM 093088 to E.N.G.M. and DK 046960 to R.T.K.

## Author Contributions

A.B.D., K.Z. and K.S. performed the experiments. A.B.D., K.Z. and K.S. analyzed the results. A.B.D., R.T.K. and E.N.G.M. designed the experiments. A.B.D. and E.N.G.M. wrote the paper.

## SUPPLEMENTARY INFORMATION

### Conditions for LC-MS analysis of 5’-deoxyadenosine

Benzoyl chloride-derivatized samples were analyzed in triplicate by UHPLC-tandem mass spectrometry using a Phenomenex Kinetex C18 HPLC column (2.1 mm × 100 mm, 1.7 μm) on a Thermo Fisher Vanquish UHPLC system interfaced to a Thermo Fisher TSQ Quantum Ultra triplequadrupole mass spectrometer. The injection volume was 5 μL. Mobile phase A consisted of 10 mM ammonium formate with 0.15% formic acid. Mobile phase B was pure acetonitrile. The flow rate was 600 μL/min, and the gradient was as follows: initial, 5% B; 0.01 min, 8% B; 0.60 min, 26% B; 0.68 min, 75% B; 1.06 min, 100% B; 1.81 min, 100% B; 2.18 min, 0% B; 2.28 min, 0% B; 3.00 min. Positive electrospray ionization mode was used, with the capillary set at 2.5 kV. The capillary temperature was 400 °C, vaporizer temperature was 400 °C, aux gas pressure was 5, and sheath gas pressure was 10. Detection was performed in dynamic MRM modeusing the conditions listed in Table S3. Peaks were integrated using Thermo Xcalibur QuanBrowser. All peaks were inspected to ensure correct integration.

**Figure S1:**
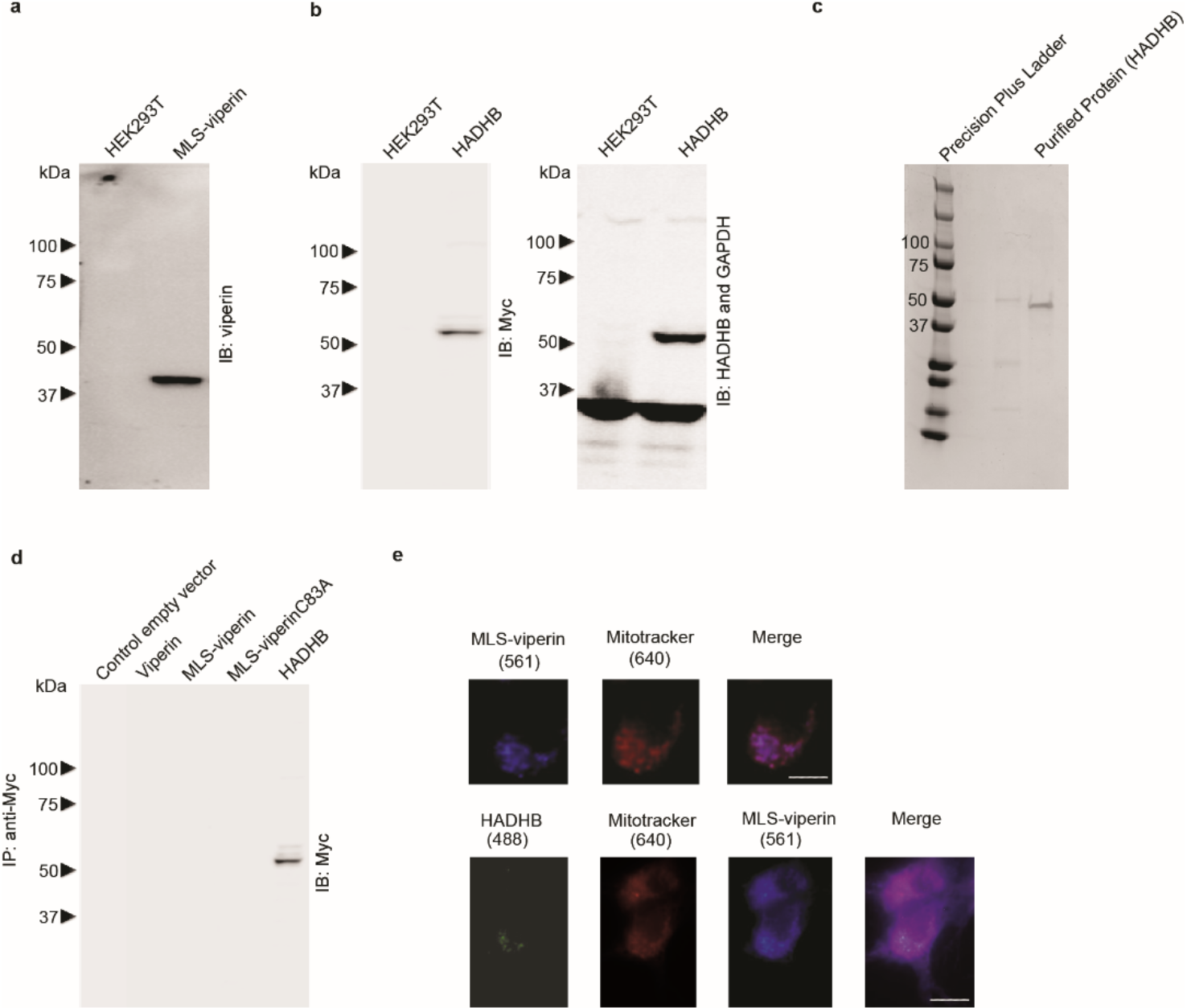
**a, b** Validation of antibodies used for immunoblotting of MLS-viperin and HADHB. Lysates from non-transfected cells or from transiently transfected cell lines expressing only MLS-viperin or HADHB were immunoblotted, probed with antibodies as indicated on the figure panels and subsequently visualized with peroxidase-labeled goat anti-rabbit IgG or goat anti-mouse IgG antibodies. **c** Coomassie-stained gel for the recombinant human HADHB protein purchased from AdooQ^®^ bioscience. **d** Specificity of antibodies used in pull-down experiments. No cross-reactivity was observed of the anti-Myc antibody used to immuno-tag HADHB with WT-viperin, MLS-viperin, and MLS-viperinC83A. Whole-cell extracts from HEK 293T were subjected to immunoprecipitation and immunoblotting with the indicated antibodies. **e** MLS-viperin localized to mitochondrion and colocalized with HADHB. HEK 293T cells were transiently transfected with MLS-viperin and/or HADHB. 14 h post-transfection mitochondrial staining was done using MitoTracker™ Deep Red FM (M22426, Thermo Scientific) as recommended by the manufacturer. After washing off the excess mitochondrial stain, cells were fixed in 4% paraformaldehyde, permeabilized with 0.05% Triton-X, and immuno-stained for viperin and HADHB. Images shown are representative of n = 10 cells. Scale bar = 5 μm. *Top panels:* cells transfected with MLS-viperin (purple) and mitochondrion stained with MitoTracker (red) showed colocalization at the mitochondrion. *Bottom panels*: cells were cotransfected with MLS-viperin and HADHB. Staining for Mitochondrion (red), MLS-viperin (purple) and HADHB (green) demonstrate co-localization of MLS-viperin and HADHB at the mitochondrion.

**Figure S2:**
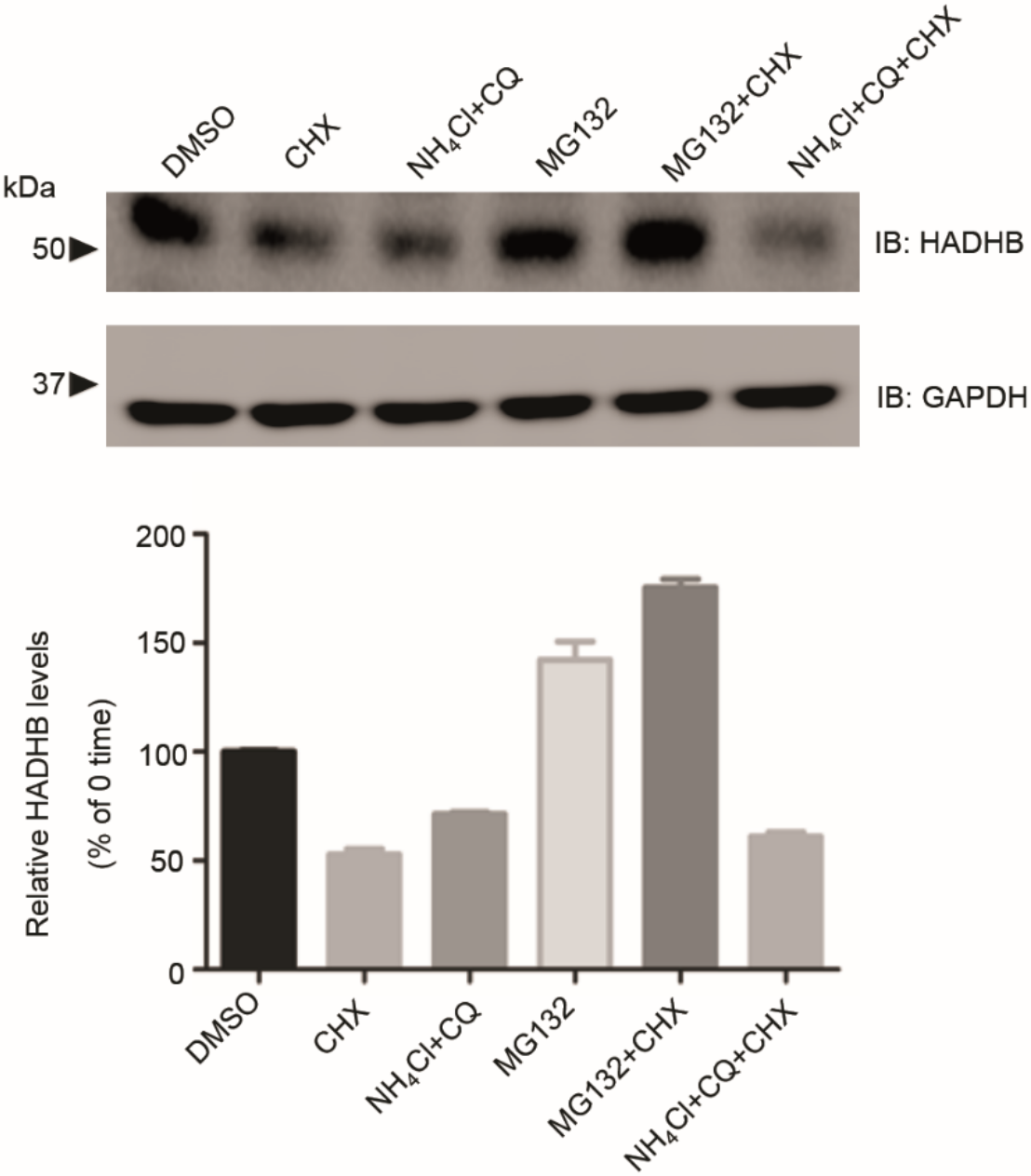
Pathway of HADHB degradation. HEK 293T cells transfected with Myc-HADHB for 24 h were treated with or without cycloheximide (20 μM) and either the proteasomal inhibitor MG132 (1 μM) or the lysosomal inhibitors ammonium chloride (50 μM) and chloroquine (50 μM) for 12 h. Levels of HADHB were then determined by immunoblot analysis and quantified by densitometry. Data are presented as mean ± SE, n = 6. GAPDH served as a loading control in all experiments. Data indicate that HADHB is retro-translocated from the mitochondrion and degraded by the proteasomal pathway.

**Figure S3:**
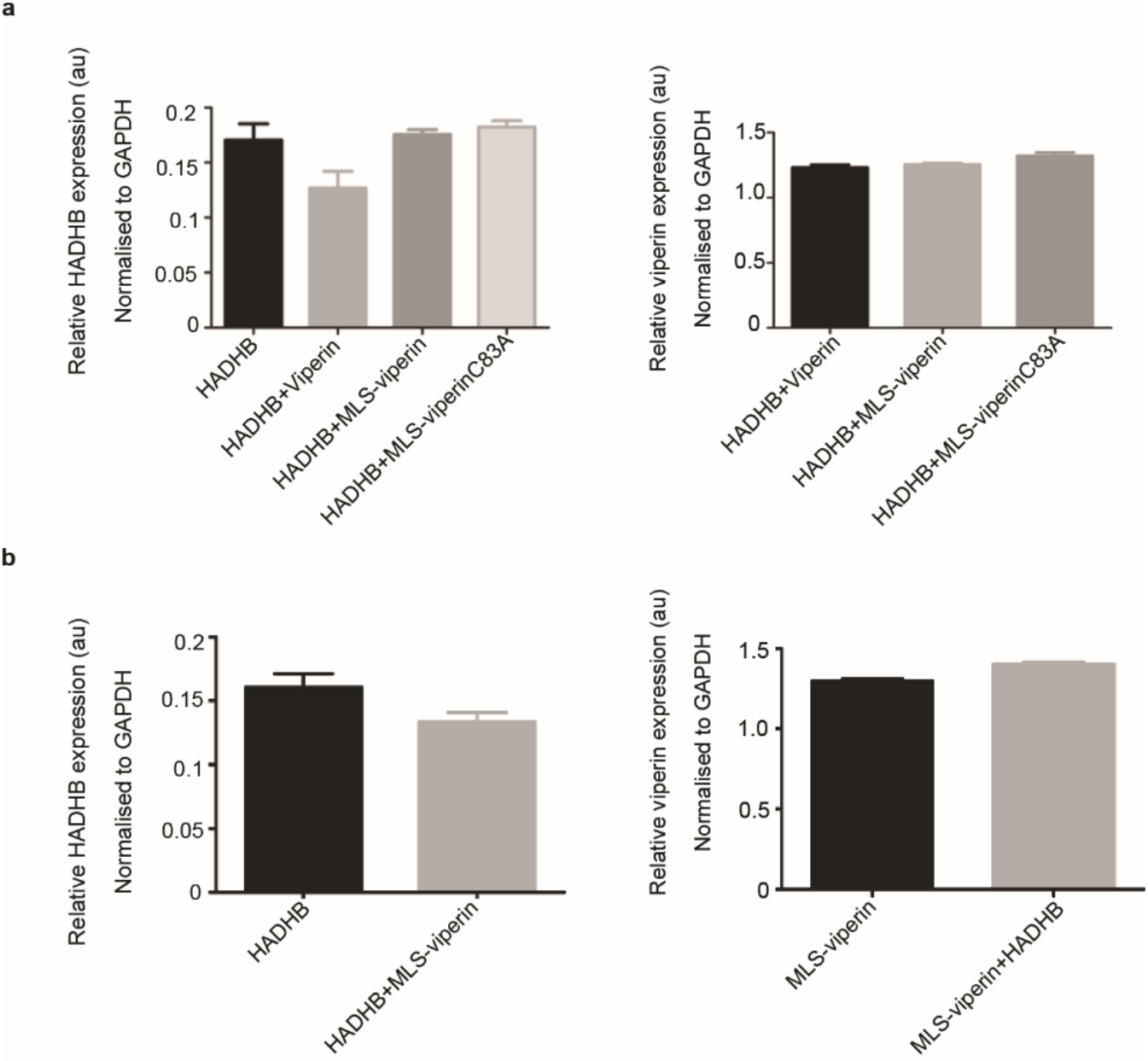
Analysis of the expression levels of HADHB, MLS-viperin, and MLS-viperinC83A in transfected HEK 293T lysates used for enzyme assays, as discussed in the main text. HEK 293T cells were transfected with the indicated genes and cell lysates subjected to immunoblot analysis using antibodies as indicated in Fig. 4 of the main text. The levels of HADHB, MLS-viperin, and MLS-viperinC83A were then determined by immunoblot analysis and quantified by densitometry. Data are presented as mean ± SE, n = 6. GAPDH served as a loading control in all experiments.

**Table S1:** Apparent *k*_cat_ (h^-1^) for ΔN-viperin under various conditions. Activity determined by the amount of 5’-dA produced in 1 h under the assay conditions.

| Condition | Turnovers ($\text{h}^{-1}$) |
| --- | --- |
| $\Delta\text{N}$-viperin | $0.51 \pm 0.08$ |
| $\Delta\text{N}$-viperin + CTP | $1.75 \pm 0.03$ |
| $\Delta\text{N}$-viperin + HADHB | $2.23 \pm 0.04$ |
| $\Delta\text{N}$-viperin + HADHB + CTP | $4.44 \pm 0.06$ |

**Table S2:** Apparent *k*_cat_ (h^-1^) for MLS-viperin in lysates of HEK 293T cells under various conditions.

| Condition | Viperin (pmol) | 5'-dA (pmol) | Turnovers ($\text{h}^{-1}$) |
| --- | --- | --- | --- |
| MLS-viperin | $0.73 \pm 0.03$ | $1.14 \pm 0.02$ | $1.56 \pm 0.07$ |
| MLS-viperin + HADHB | $0.94 \pm 0.04$ | $6.31 \pm 0.04$ | $6.71 \pm 0.26$ |
| MLS-viperin+ CTP | $0.74 \pm 0.03$ | $8.97 \pm 0.05$ | $12.1 \pm 0.5$ |
| MLS-viperin + HADHB+ CTP | $0.92 \pm 0.03$ | $24.94 \pm 0.03$ | $27.1 \pm 0.9$ |

**Table S3.** MRM conditions for the target analytes

| Analyte | Precursor (m/z) | Product (m/z) | Tube Lens | Retention Time (min) |
| --- | --- | --- | --- | --- |
| **Bz-SAM** | 711.2 | 205.9 | 92 | 1.54 |
| **$^{13}\text{C}$ Bz-SAM** | 729.5 | 212.0 | 92 | 1.54 |
| **Bz-Ado** | 476.1 | 136.0 | 75 | 1.63 |
| **$^{13}\text{C}$ Bz-Ado** | 488.0 | 136.0 | 75 | 1.63 |
| **Bz-5'dA** | 460.2 | 325.0 | 80 | 1.72 |
| **$^{13}\text{C}$ Bz-5'dA** | 472.0 | 111.0 | 76 | 1.72 |

## References

1. Horner, S. M. (2014) Activation and evasion of antiviral innate immunity by hepatitis c virus. J. Mol. Biol. 426, 1198–1209

2. Horst, D., Verweij, M. C., Davison, A. J., Ressing, M. E., and Wiertz, E. (2011) Viral evasion of t cell immunity: Ancient mechanisms offering new applications. Curr. Opin. Immunol. 23, 96–103

3. Iannello, A., Debbeche, O., Martin, E., Attalah, L. H., Samarani, S., and Ahmad, A. (2006) Viral strategies for evading antiviral cellular immune responses of the host. J. Leukoc. Biol. 79, 16–35

4. Jiang, X. M., and Chen, Z. J. J. (2012) The role of ubiquitylation in immune defence and pathogen evasion. Nat. Rev. Immunol. 12, 35–48

5. Taylor, K. E., and Mossman, K. L. (2013) Recent advances in understanding viral evasion of type i interferon. Immunology 138, 190–197

6. Bhoj, V. G., and Chen, Z. J. (2009) Ubiquitylation in innate and adaptive immunity. Nature 458, 430–437

7. Jo, E. K., Yang, C. S., Choi, C. H., and Harding, C. V. (2007) Intracellular signalling cascades regulating innate immune responses to mycobacteria: Branching out from tolllike receptors. Cell Microbiol. 9, 1087–1098

8. O’Neill, L. A., and Bowie, A. G. (2010) Sensing and signaling in antiviral innate immunity. Curr Biol 20, R328–333

9. Schneider, W. M., Chevillotte, M. D., and Rice, C. M. (2014) Interferon-stimulated genes: A complex web of host defenses. in Ann. Rev. Immunol. 32, 513–545

10. Morrison, J., Aguirre, S., and Fernandez-Sesma, A. (2012) Innate immunity evasion by dengue virus. Viruses-Basel 4, 397–413

11. Orange, J. S., Fassett, M. S., Koopman, L. A., Boyson, J. E., and Strominger, J. L. (2002) Viral evasion of natural killer cells. Nat. Immunol. 3, 1006–1012

12. Chin, K. C., and Cresswell, P. (2001) Viperin (cig5), an ifn-inducible antiviral protein directly induced by human cytomegalovirus. Proc. Natl. Acad. Sc.i U. S.A. 98, 15125–15130

13. Seo, J. Y., Yaneva, R., Hinson, E. R., and Cresswell, P. (2011) Human cytomegalovirus directly induces the antiviral protein viperin to enhance infectivity. Science 332, 1093–1097

14. Helbig, K. J., and Beard, M. R. (2014) The role of viperin in the innate antiviral response. J. Mol. Biol. 426, 1210–1219

15. Mattijssen, S., and Pruijn, G. J. (2012) Viperin, a key player in the antiviral response. Microbes Infect. 14, 419–426

16. Chakravarti, A., Selvadurai, K., Shahoei, R., Lee, H., Fatma, S., Tajkhorshid, E., and Huang, R. H. (2018) Reconstitution and substrate specificity for isopentenyl pyrophosphate of the antiviral radical SAM enzyme viperin. J. Biol. Chem. 293, 14122–14133

17. Honarmand Ebrahimi, K., Carr, S. B., McCullagh, J., Wickens, J., Rees, N. H., Cantley, J., and Armstrong, F. A. (2017) The radical SAM enzyme viperin catalyzes reductive addition of a 5’-deoxyadenosyl radical to udp-glucose in vitro. FEBS Lett. 591, 2394–2405

18. Duschene, K. S., and Broderick, J. B. (2010) The antiviral protein viperin is a radical sam enzyme. FEBS Lett. 584, 1263–1267

19. Landgraf, B. J., McCarthy, E. L., and Booker, S. J. (2016) Radical s-adenosylmethionine enzymes in human health and disease. Annu. Rev. Biochem. 85, 485–514

20. Wang, J., Woldring, R. P., Roman-Melendez, G. D., McClain, A. M., Alzua, B. R., and Marsh, E. N.G. (2014) Recent advances in radical sam enzymology: New structures and mechanisms. ACS Chem. Biol. 9, 1929–1938

21. Duschene, K. S., and Broderick, J. B. (2012) Viperin: A radical response to viral infection. Biomol. Concepts 3, 255–266

22. Vey, J. L., and Drennan, C. L. (2011) Structural insights into radical generation by the radical sam superfamily. Chem. Rev. 111, 2487–2506

23. Marsh, E. N.G., Patterson, D. P., and Li, L. (2010) Adenosyl radical: Reagent and catalyst in enzyme reactions. ChemBioChem 11, 604–621

24. Frey, P. A., Hegeman, A. D., and Ruzicka, F. J. (2008) The radical SAM superfamily. Crit Rev. Biochem. Mol. Biol. 43, 63–88

25. Marsh, E. N.G, Patwardhan, A., and Huhta, M. S. (2004) S-adenosylmethionine radical enzymes. Bioorg. Chem. 32, 326–340

26. Cheek, J., and Broderick, J. B. (2001) Adenosylmethionine-dependent iron-sulfur enzymes: Versatile clusters in a radical new role. J. Biol. Inorg. Chem. 6, 209–226

27. Gizzi, A. S., Grove, T. L., Arnold, J. J., Jose, J., Jangra, R. K., Garforth, S. J., Du, Q., Cahill, S. M., Dulyaninova, N. G., Love, J. D., Chandran, K., Bresnick, A. R., Cameron, C. E., and Almo, S. C. (2018) A naturally occurring antiviral ribonucleotide encoded by the human genome. Nature 558, 610–614

28. Saitoh, T., Satoh, T., Yamamoto, N., Uematsu, S., Takeuchi, O., Kawai, T., and Akira, S. (2011) Antiviral protein viperin promotes toll-like receptor 7- and toll-like receptor 9-mediated type i interferon production in plasmacytoid dendritic cells. Immunity 34, 352–363

29. Makins, C., Ghosh, S., Roman-Melendez, G. D., Malec, P. A., Kennedy, R. T., and Marsh, E. N.G. (2016) Does viperin function as a radical s-adenosyl-l-methionine-dependent enzyme in regulating farnesylpyrophosphate synthase expression and activity? J. Biol. Chem. 291, 26806–26815

30. Seo, J. Y., and Cresswell, P. (2013) Viperin regulates cellular lipid metabolism during human cytomegalovirus infection. PLoS Pathog. 9, e1003497

31. Wang, X., Hinson, E. R., and Cresswell, P. (2007) The interferon-inducible protein viperin inhibits influenza virus release by perturbing lipid rafts. Cell Host Microbe 2, 96–105

32. Eom, J. et. al. (2019) Intrinsic expression of viperin regulates thermogenesis in adipose tissues. Proc. Natl. Acad. Sci. (USA) Early view

33. Dumbrepatil, A. B., Ghosh, S., Zegalia, K. A., Malec, P. A., Hoff, J. D., Kennedy, R. T., and Marsh, E. N.G. (2019) Viperin interacts with the kinase IRAK1 and the E3 ubiquitin ligase TRAF6, coupling innate immune signaling to antiviral ribonucleotide synthesis. J. Biol. Chem. 294, 6888–6898

34. Hee, J. S., and Cresswell, P. (2017) Viperin interaction with mitochondrial antiviral signaling protein (mavs) limits viperin-mediated inhibition of the interferon response in macrophages. PLoS One 12, e0172236

35. Bartlett, K., and Eaton, S. (2004) Mitochondrial beta-oxidation. Eur. J. Biochem. 271, 462–469

36. Zhou, Z. Q., Zhou, J. H., and Du, Y. C. (2012) Estrogen receptor beta interacts and colocalizes with hadhb in mitochondria. Biochem. Biophys. Res. Comm. 427, 305–308

37. Panayiotou, C., Lindqvist, R., Kurhade, C., Vonderstein, K., Pasto, J., Edlund, K., Upadhyay, A. S., and Overby, A. K. (2018) Viperin restricts zika virus and tick-borne encephalitis virus replication by targeting NS3 for proteasomal degradation. J. Virol. 92, e02054–02017

38. Pickles, S., Vigie, P., and Youle, R. J. (2018) Mitophagy and quality control mechanisms in mitochondrial maintenance. Curr. Biol. 28, R170–R185

39. Taylor, E. B., and Rutter, J. (2011) Mitochondrial quality control by the ubiquitin-proteasome system. Biochem. Soc. Trans. 39, 1509–1513

40. Schwinghammer, K., Cheung, A. C. M., Morozov, Y. I., Agaronyan, K., Temiakov, D., and Cramer, P. (2013) Structure of human mitochondrial RNA polymerase elongation complex. Nat. Struct. Molec. Biol. 20, 1298–U1225

41. Wong, J. M. T., Malec, P. A., Mabrouk, O. S., Ro, J., Dus, M., and Kennedy, R. T. (2016) Benzoyl chloride derivatization with liquid chromatography-mass spectrometry for targeted metabolomics of neurochemicals in biological samples. J. Chromatog. A 1446, 78–90

